# ProDualNet: Dual-Target Protein Sequence Design Method Based on Protein Language Model and Structure Model

**DOI:** 10.1101/2025.02.28.640919

**Authors:** Liu Cheng, Ting Wei, Xiaochen Cui, Haifeng Chen, Zhangsheng Yu

**Affiliations:** Department of Bioinformatics and Biostatistics, School of Life Sciences and Biotechnology, Shanghai Jiao Tong University, Shanghai, China; Intelligent Medicine Original (Shanghai) Co., Ltd., Shanghai, China; SJTU-Yale Joint Center for Biostatistics and Data Science Organization, Shanghai Jiao Tong University, Shanghai, China; Center for Biomedical Data Science, Translational Science Institute, Shanghai Jiao Tong University School of Medicine, Shanghai, China; Clinical Research Institute, Shanghai Jiao Tong University School of Medicine, Shanghai, China

## Abstract

Proteins typically interact with multiple partners to regulate biological processes, and peptide drugs targeting multiple receptors have shown strong therapeutic potential, emphasizing the need for multi-target strategies in protein design. However, most current protein sequence design methods focus on interactions with a single receptor, often neglecting the complexity of designing proteins that can bind to two distinct receptors. We introduced ProDualNet, a novel approach for designing dual-target protein sequences by integrating sequence-structure information from two distinct receptors. ProDualNet used a heterogeneous graph network for pretraining and combines noise-augmented single-target data with real dual-target data for fine-tuning. This approach addressed the challenge of limited dual-target protein experimental structures. The efficacy of ProDualNet has been validated across multiple test sets, demonstrating better recovery and success rates compared to other multi-state design methods. ***In silico*** evaluation of cases like dual-target allosteric binding and non-overlapping interface binding highlights its potential for designing dual-target binding proteins. Furthermore, we validated ProDualNet’s ability to model the relationships between sequences, structures, and functions by zero-shot prediction tasks, including dual-target protein functional effects and mutant functional effects.

## Introduction

In living organisms, proteins rarely function in isolation. Instead, they engage in intricate networks of interactions with other proteins, essential for cellular homeostasis and regulating physiological processes^1–3^. A single protein often interacts with multiple distinct partners, forming complexes that mediate a broad range of biological functions. The phenomenon of multi-target binding exemplifies the complexity of biological signaling networks^4,5^. For instance, Tirzepatide, a co-agonist peptide analog, simultaneously binds to both GIP and GLP-1 receptors, harnessing their synergistic effects to improve blood glucose and body weight regulation, offering superior effects over monotherapies targeting either receptor alone^6,7^. This dual-receptor activation highlights the potential of multi-target drug design as a promising therapeutic strategy^8–14^.

Recent advances in AI-driven design tools^15–18^, such as ProteinMPNN^19^ and ESM-IF1^20^, have transformed protein sequence design. Known as protein inverse folding, this approach aims to generate protein sequences that fold into specific protein structures and possess desired functions^21^, starting from a single protein backbone structure.

While some methods focus on multi-conformation design^20,22–24^, most protein sequence design research remains centered on single-target proteins, typically using the structure of a single protein complex^25^. This approach neglects the intricate interactions between the designed protein and multiple target proteins^1,8^, focusing primarily on binding to only one target. In many multi-state design methods, the probabilities of different conformations are simply averaged, without fully exploiting the model to explore the interactions between the designed protein and multiple receptors. As a result, these methods fail to capture the varying binding information when the protein interacts with different receptors. Therefore, multi-target design requires integrating multiple protein complex structures, ensuring that the designed protein can effectively bind to different receptors and perform its intended function. A further challenge lies in the limited availability of experimental datasets for multi-target protein complexes, which significantly hinders effective training.

To address these challenges, we introduce the Protein Dual-Target Design Network (ProDualNet), a novel model for designing dual-target protein sequences. ProDualNet improves the model to design proteins that interact with two targets by jointly modeling the two conformations of dual-target proteins. We first developed a protein sequence design framework based on heterogeneous graph networks, which serves as a pretraining model. This framework excels in capturing geometric features, which are crucial for accurately modeling the relationships between protein sequences, structures, and functions. To overcome the scarcity of dual-target experimental structures, we employed a noise-augmentation strategy (NoiseMix) during fine-tuning, which combines augmented single-target data with experimental dual-target data. This approach improves the model to capture the joint distributions of multiple protein complexes, which is essential for effective dual-target protein design.

We applied ProDualNet to various design tasks, showing its robustness and effectiveness. Additionally, we assessed its performance as a zero-shot predictor on dual-target sequence functional evaluations and mutation datasets, proving its capability to learn complex relationships between protein sequences, structures, and functions^26,27^.

## Results

### Model Architecture

ProDualNet is a dual-target protein sequence design framework that integrates the structure and sequence information of two protein complexes. The model is first pre-trained on single-target data (Fig. 1a) and subsequently fine-tuned on dual-target data using a noise-augmentation approach (NoiseMix), addressing the challenge posed by the limited availability of dual-target complexes in the Protein Data Bank (PDB), and enhancing training robustness (Fig. 1b).

**Figure 1:**
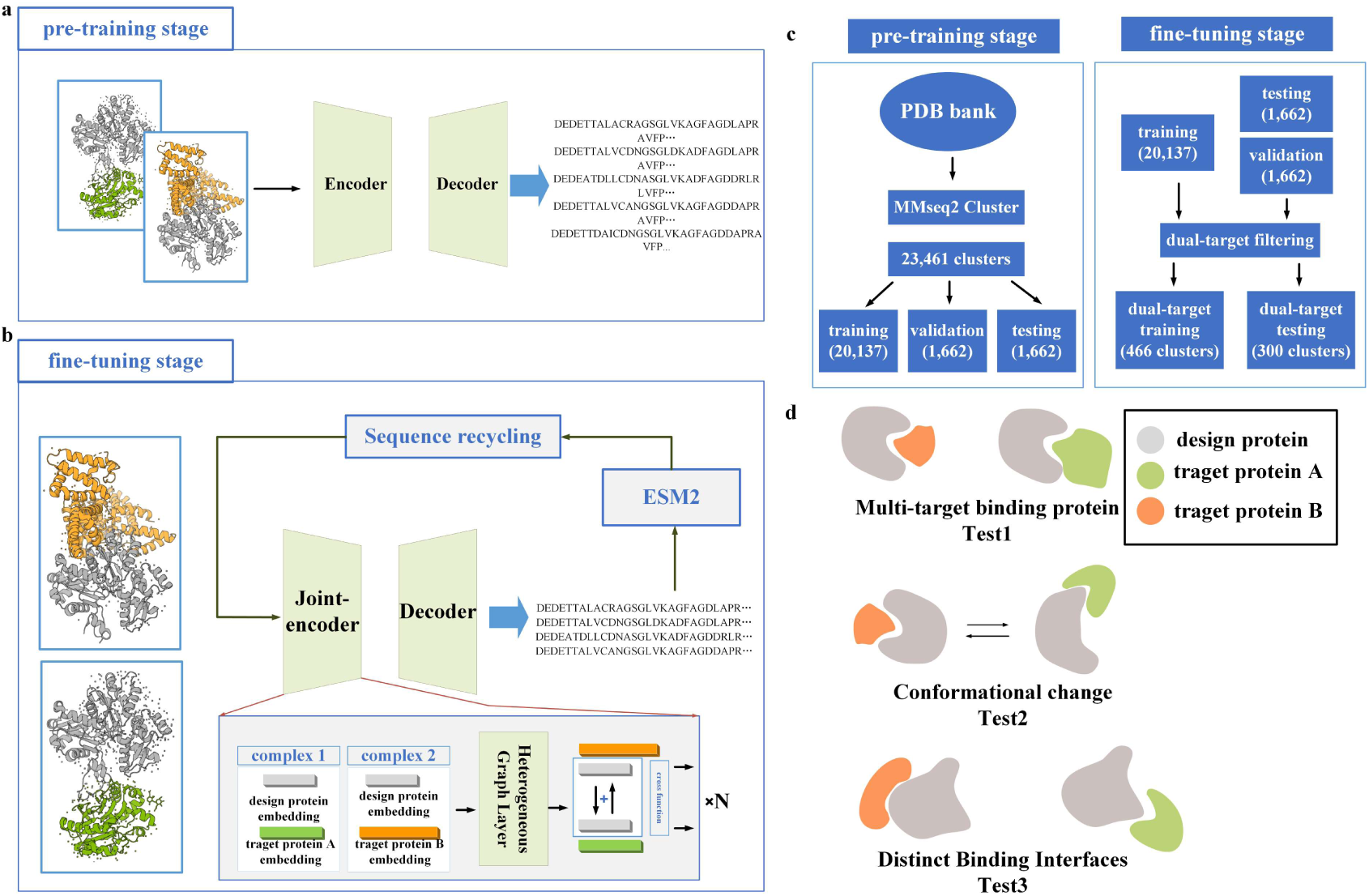
Architecture of ProDualNet. (a) Schematic representation of ProDualNet pretraining using a standard training strategy. (b) ProDualNet fine-tuning for dual-target tasks, employing a shared encoder-decoder structure and an ESM2-based recycling strategy. (c) Overview of the ProDualNet dataset construction process, comprising pretraining dataset and fine-tuning dataset. (d) Three test sets based on different evaluation criteria.

During fine-tuning, ProDualNet adopts a shared encoder-decoder architecture. The encoder, comprising four layers of heterogeneous graph networks, simultaneously processes both sequence and structural data from the dual-target complex, learning the inter-protein relationships within a unified embedding space. The heterogeneous graph network combines a Message Passing Neural Network (MPNN) with a global aggregation network. And the model employs an autoregressive decoder to predict amino acid sequences, while ESM2^28^ is incorporated to provide additional evolutionary insights (Fig. 1b). This architecture enhances the understanding of the relationship between protein design and functionality.

In the evaluation of ProDualNet, the test dataset is completely independent from the training dataset, even during the pretraining phase, effectively eliminating the risk of data leakage (Fig. 1c). In contrast, the training dataset for the comparative method, ProteinMPNN_mean, may overlap with the test data. ProteinMPNN_mean, based on a multi-state design framework, is a leading approach in the field of multi-state protein design and has been successfully applied to protein switches design task^19,24^.

### Evaluating ProDualNet on dual-target protein testing sets

To evaluate the model’s robustness under different conditions, we established three test sets based on distinct criteria: Test Set 1 focused on typical dual-target design tasks, Test Set 2 involved proteins that undergo conformational changes upon binding to different receptors, and Test Set 3 examined proteins with non-overlapping binding interfaces to two target proteins (Fig. 1d; see “Methods”). Model performance was evaluated using three primary metrics: global residue recovery, interface residue recovery, and perplexity.

Firstly, we evaluated the model’s performance on typical dual-target design tasks using Test Set 1, which consists of 159 protein complex pairs. ProDualNet, using an unconditional sampling strategy for protein complex sequence design, outperformed all comparison methods, achieving a sequence recovery of 0.581 and an interface recovery of 0.603. These results were significantly better than ProteinMPNN_mean, which had a sequence recovery of 0.517 and an interface recovery of 0.523 (Fig. 2a). Statistical analysis, including p-values, further confirmed the significance and robustness of these improvements. Specifically, in terms of perplexity, ProDualNet achieved a score of 3.9, outperforming the ProteinMPNN_mean (4.8) and ProteinMPNN_single (5.1).

**Figure 2:**
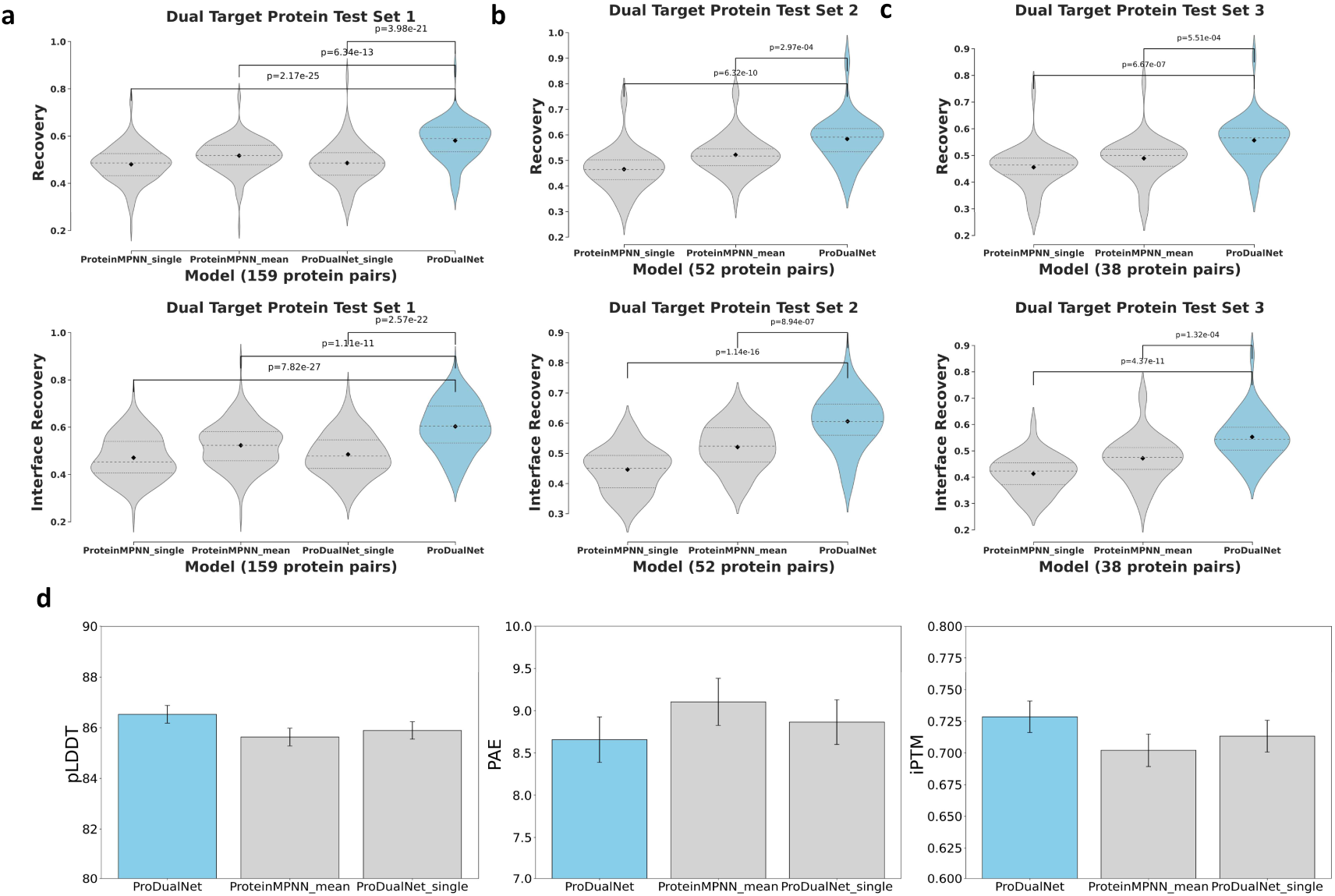
Evaluation Results in Different Design Tasks. (a) Global recovery and interface recovery on Test Set 1. (b) Evaluation results on Test Set 2, featuring conformational changes. (c) Evaluation results on Test Set 3, with varying interface combinations. (d) AlphaFold3 results for the designed sequence on Test Set 1.

Additionally, we employed AlphaFold3^29^, the state-of-the-art model for protein complex structure prediction, to predict the structures of the designed sequences and target receptors. We focus primarily on three rank metrics: pLDDT, PAE, and ipTM. These metrics provide indicators for assessing the quality of sequence designs^30,31^: higher pLDDT scores reflect more reliable local structures, higher ipTM values suggest better alignment of interface topologies, and lower PAE values indicate smaller discrepancies between predicted and actual protein structures. ProDualNet outperforms the comparison methods in terms of the overall average values of these three metrics across 159 designed protein complex pairs. Specifically, ProDualNet achieves an ipTM of 0.728, a PAE of 8.65, and a pLDDT of 86.52, thereby highlighting the reliability of the designs generated by ProDualNet (Fig. 2d).

To further assess and validate the effectiveness of our model in the dual-target protein design task, we evaluated our model under two distinct design categories, using Test Sets 2 and 3. The first category involves a conformational change in the designed protein upon binding to different receptors, comprising 52 protein complex pairs (Test Set 2; Fig. 1d). The second category examines the designed protein binding to two target receptors at distinct interfaces, consisting of 38 protein complex pairs (Test Set 3; Fig. 1d).

In the experiments with Test Set 2, ProDualNet was applied to design sequences for proteins that undergo conformational changes upon binding to different target proteins. ProDualNet achieved a sequence recovery of 0.584 and an interface recovery of 0.606, while ProteinMPNN_mean had a sequence recovery of 0.521 and an interface recovery of 0.520. ProDualNet outperformed the comparison methods in both total residue and interface residue recovery, highlighting the effectiveness of its joint modeling strategy in dual-target conformation design (Fig. 2b). In addition, AlphaFold2 shows that our design sequences transition from alpha-helix to beta-sheet upon binding to different receptors (Fig. S7).

In Test Set 3, the designed proteins had non-overlapping binding interfaces with two target proteins. We evaluated total residue recovery and interface residue recovery, and the results showed that ProDualNet outperformed the comparison methods in all metrics, particularly in interface residue recovery (Fig. 2c). Specifically, ProDualNet achieved a sequence recovery of 0.556 and an interface recovery of 0.552, while ProteinMPNN_mean reached a sequence recovery of 0.489 and an interface recovery of 0.472. Furthermore, when ProteinMPNN_single was used with single-complex models, both metrics were significantly lower, with a sequence recovery of 0.455 and an interface recovery of 0.412.

To further assess the design capabilities of ProDualNet, we adopted an autoregressive strategy, designing 10 sequences for each of the 38 cases in Test Set 3, followed by rapid structure predictions using AlphaFold2^32,33^. For comparison, we selected two variants of ProteinMPNN: ProteinMPNN_mean and ProteinMPNN_b (ProteinMPNN single best). ProteinMPNN_b generates sequences independently for each target protein complex, neglecting the interrelationship between distinct target proteins in dual-target design tasks and the conformational changes of the designed protein across different states. We used PAE and pLDDT as evaluation metrics to assess both the accuracy of the predicted structures and the reliability of the designed sequences^30^.

As shown in Fig. 3a, for the 38 cases in Test Set 3, ProDualNet outperformed ProteinMPNN_mean in 20 tasks, with a pLDDT of 88.37 and a PAE of 10.79, compared to ProteinMPNN_mean’s PLDDT of 87.86 and PAE of 11.49. Furthermore, ProDualNet demonstrated comparable or superior performance to ProteinMPNN_b across the majority of tasks. These results suggest that ProDualNet is more effective than single-complex-based design methods in dual-target protein design. This improvement is likely due to the additional conformation information provided when the designed protein binds to two distinct targets, enabling ProDualNet to better map the protein sequence space to structure space. This highlights the advantages of the multi-conformation information in protein sequence design^22,23^.

**Figure 3:**
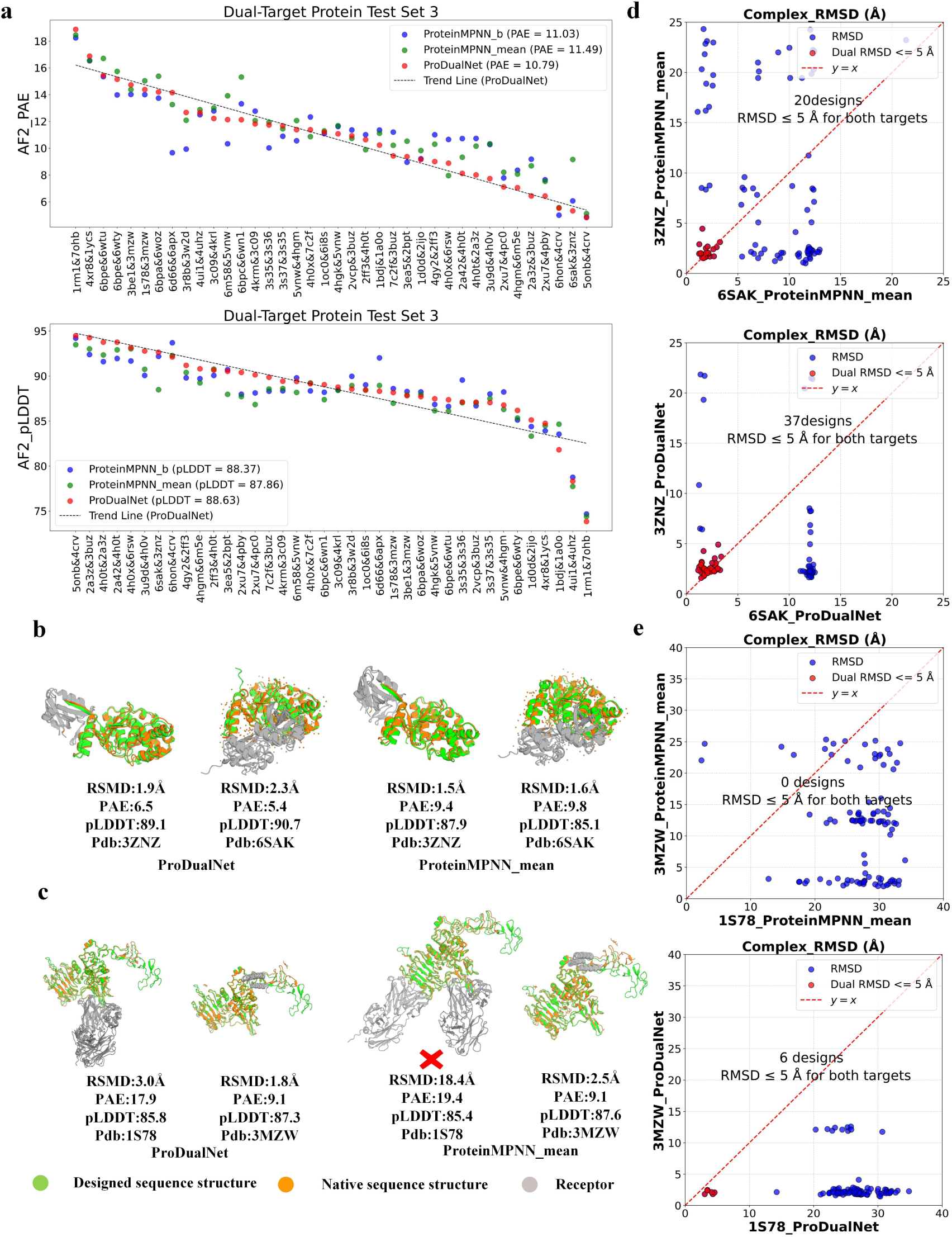
Comparison of Protein Design Methods Across Different Design Cases. (a) Comparison of AlphaFold2 scores (PAE, pLDDT) between ProteinMPNN_b (ProteinMPNN_single_best), ProteinMPNN_mean, and ProDualNet across 38 design tasks in Test Set 3. (b) AlphaFold3 prediction results for successful and failed designs: ProDualNet vs. ProteinMPNN_mean in the Ubiquitin thioesterase Otulin case (275 amino acids, PDB id: 3ZNZ, 6SAK). (c) erbB-2 receptor design case (624 amino acids, PDB id: 1S78, 3MZW): Green represents the designed structure, yellow represents the native structure, and gray represents the receptor. (d) Comparison of complex RMSD and success rate in the Ubiquitin thioesterase Otulin (length, 275 amino acids) design case: ProDualNet vs. ProteinMPNN_mean. The x-axis and y-axis represent protein complex RMSD values between the designed sequences and two different receptors, compared to the native structure. (e) Comparison of complex RMSD and success rate in the erbB-2 receptor (length, 624 amino acids) design case.

### Case Studies in Dual-Target Protein Design

To further validate the effectiveness of our approach, we selected two design cases from Test Set 3: Ubiquitin thioesterase otulin and Receptor protein-tyrosine kinase erbB-2, which bind to different receptors at distinct interfaces. For these cases, we designed 100 sequences using both ProteinMPNN_mean and ProDualNet, and predicted the protein complex structures of designed sequence using AlphaFold3.

The success rate of each design task was determined by whether the designed proteins could correctly bind within the interaction space of the target receptors. We calculated the Root Mean Square Deviation (RMSD) of the protein complexes, defining a successful design as having an RMSD ≤ 5 Å for both protein complexes ^31^.

Specifically, in the Ubiquitin thioesterase otulin design task (PDB id: 3ZNZ, 6SAK), the designed protein sequence consisted of 275 amino acids. ProDualNet achieved a success rate of 37%, compared to 20% for ProteinMPNN_mean (Fig. 3d). Furthermore, ProDualNet outperformed ProteinMPNN_mean in AF3 metrics such as PAE and pLDDT (Fig. 3b, Fig. S10 and Fig. S11a), see supplementary materials.

In the more challenging Receptor protein-tyrosine kinase erbB-2 design task (PDB id: 1S78, 3MZW), where the designed protein sequence contained 624 amino acids, ProteinMPNN_mean failed to design any acceptable sequences (Fig. 3c and Fig. 3e). In contrast, ProDualNet successfully produced 6 high-quality designs out of 100, and outperformed ProteinMPNN_mean in AF3 metrics such as PAE and pLDDT (Fig. S11b) proving its robust design capabilities.

A detailed analysis of AlphaFold3 predictions revealed that while the protein designed by ProteinMPNN_mean could bind to Immunoglobulin G-binding protein A (PDB id: 3MZW), while incorrect binding interfaces for interactions with Pertuzumab Fab light chain and Pertuzumab Fab heavy chain (PDB id: 1S78, Fig. 3c).

These results indicate that ProDualNet, by jointly modeling dual-target data, achieves higher accuracy and success rates in more complex and challenging design tasks compared to ProteinMPNN_mean.

To further investigate the advantages of dual-target protein design, we conducted experiments based on single-target protein complex design. In this experiment, we employed ProteinMPNN’s single-target design method (ProteinMPNN_a) and directly predicted the protein complex structures for the designed sequences with another receptor using AlphaFold3. The results show that both ProDualNet and ProteinMPNN_mean outperformed the single-conformation method (Fig. 4a). The failure of the single-target approach can be attributed to its inability to get the binding interface information between the designed protein and different target receptors.

**Figure 4:**
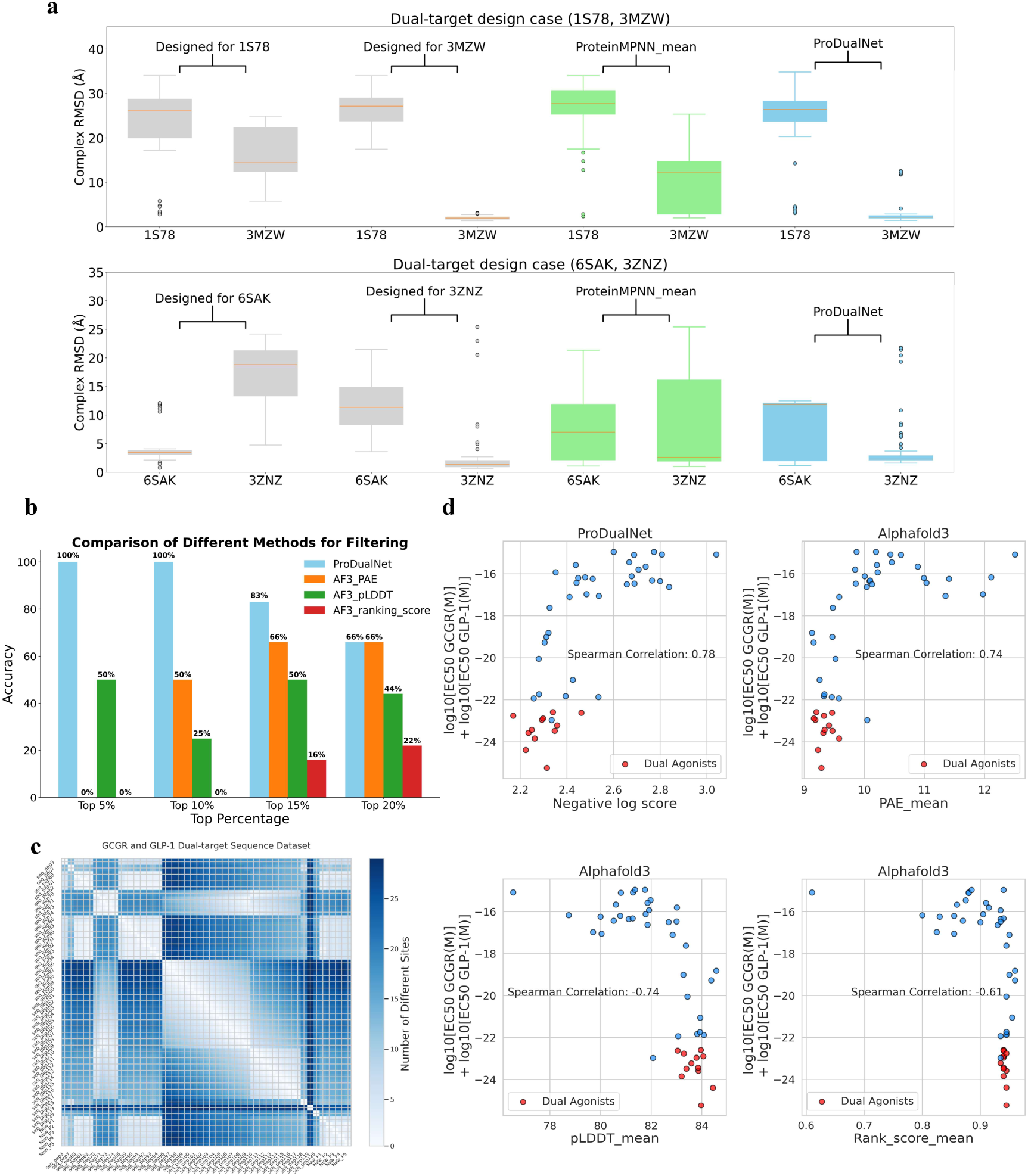
Evaluation of ProDualNet’s Advantage in Sequence Design and Functional Scoring for Dual-Target Agonist. (a) AlphaFold3 structure prediction results for sequences designed for single target structure and directly predicted for another. (b) Comparative analysis of positive dual-target sequences using the top-k% selection strategy to assess the model’s ability to predict functionality. (c) Sequence differences in the GCGR/GLP-1R dual-target test set. (d) Comparison of ProDualNet dual agonist sequence rank scores with AlphaFold3 rank scores. Red dots represent dual agonists, while blue dots represent non-dual agonists.

### Functional Effect Prediction for Dual-Target Sequences

Accurately explaining the functional changes induced by protein mutations or redesigned protein sequences is a key aspect of both directed evolution and AI-driven protein design^26,27,34,35^. A model’s ability to predict the effects of amino acid changes reflects its understanding of the relationship between sequence, structure, and function, which in turn determines its capacity to generate sequences with specific functionalities. Since zero-shot screening often relies on Negative Log-Likelihood (NLL), this score reflects the model’s preference for generating output sequences. In addition, by exploring the use of sequence design models for zero-shot functional predictions, we can enable rapid ***in silico*** screening of candidate sequences, ultimately improving the efficiency of subsequent wet-lab experiments^34,35^.

In this study, we evaluated the potential of ProDualNet for dual-target peptide-based drug design using a benchmark dataset of dual-target agonists derived from the GCGR and GLP-1R receptors. The dataset consists of 46 protein sequences, each comprising 29 amino acids, with experimental EC50 data reflecting their interactions with both the GCGR and GLP-1R receptors^37^. A detailed distribution of the dataset is shown in the Figure 4(c). This task considerably more challenging than single-site mutation prediction. Importantly, none of the sequences in the test set were part of the model’s training data, ensuring a robust evaluation. Additionally, ESM2 features were excluded from the analysis.

To assess the zero-shot prediction capability of ProDualNet, we utilized the Negative Log-Likelihood (NLL) scores as a unified ranking metric for the binding affinity between designed sequences and two target receptors^38^. Specifically, lower NLL scores were strongly correlated with a higher probability of successful binding to both receptors. Experimental results revealed a Spearman correlation coefficient of 0.78 between the NLL scores and the combined log10EC50 values for the two receptors, significantly outperforming the results obtained with AlphaFold3 scores (Fig. 4d). The higher the correlation between NLL and combined log10EC50 values, the greater the probability of producing dual agonists for our model.

To further validate the efficacy of virtual screening process, we applied the top-K% strategy, wherein sequences are ranked according to their NLL scores, and the top K% are selected for virtual experimental validation. This approach involves counting the number of sequences in the top-K% that successfully achieved dual-target design. When compared to AlphaFold3’s three scoring metrics (PAE, pLDDT, and ranking_score), ProDualNet consistently exhibited superior performance in top-K% screening (Fig. 4b). Notably, PAE and pLDDT outperformed ranking_score in sequence ranking (Fig. 4d).

### Zero-shot Prediction of Mutation Effects

To assess the ability of our pre-trained model (ProDualNet_single) to capture the relationships between sequence, structure, and function, we focus on mutation effect prediction. This task is critical for understanding protein function, as it directly reflects the model’s capacity to capture these intricate relationships^34,36,39^. Notably, our approach to mutation effect prediction, like the dual-target peptide functional effect prediction, does not rely on ESM2 features. This choice allows us to explore alternative modeling strategies for evaluating mutation impacts, offering a framework for optimizing protein stability and function.

We evaluated the performance of our model on two large, established benchmark datasets: the DMS stability^39,40^ and Proteingym Stability databases^27^. In the thermal stability mutation dataset from Proteingym Stability, our pretrained model achieved an average Spearman correlation coefficient of 0.67, outperforming structural pretraining-based models such as ProteinMPNN^19^ and ESM-IF1^20^ (Fig. 5a), they have been proven to be effective zero-shot likelihood models^41^. Our method also outperforms methods that integrate protein language models with structure model^42,43^. Furthermore, in the prediction of higher-order mutations, our model achieved better performance, our overall Spearman correlation coefficient is 0.45 (Fig. 5b). In the DMS stability database, which contains 298 protein systems, our model surpassed competing methods, underscoring the robustness of our approach (Fig. 5c).

**Figure 5:**
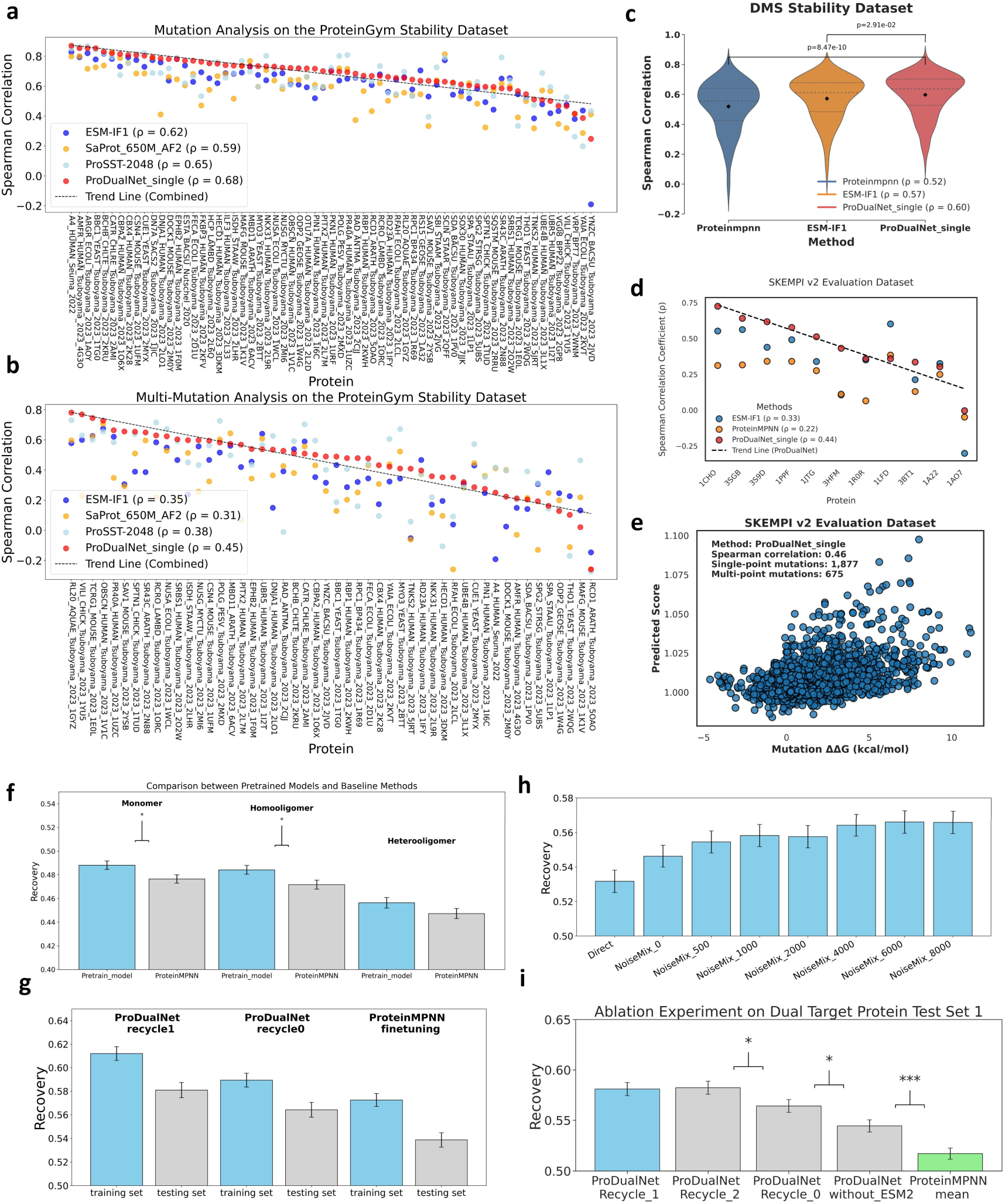
Evaluation of ProDualNet in Mutation Prediction Tasks and Ablation Studies. (a) Overall Spearman correlation coefficient evaluation of 65 sets of protein mutation data in the Proteingym Stability database, each set containing multiple mutant sequences for a protein. (b) Higher-order mutation evaluation of 65 sets of protein mutation data in the Proteingym Stability database. (c) Overall evaluation of 298 sets of protein mutation data in the DMS stability dataset. (d) Affinity evaluation of 11 sets of mutant sequences in the SKEMPI V2 database. (e) Evaluation of the relationship between model-predicted scores and binding free energy for 1877 single-point mutations and 675 multi-point mutations across 11 proteins. (f) Comparative analysis between the ProDualNet pre-trained model and ProteinMPNN. (g) Comparison of fine-tuning ProteinMPNN_mean and ProDualNet on training and test sets 1. (h) Analysis of the mixed augmented data ratio in the Noisemix strategy. (i) Evaluation of the effect of the ESM2 language model and validation of the recycling strategy.

To further assess the adaptability of our pretrained model in predicting fitness scores for complex mutations, we selected antibody mutation data from the SKEMPI V2 database^44^, focusing on samples with over 100 mutant sequences, and performed affinity scoring (Fig. 5d). When compared to other structure-based sequence design models^19,20^, our model demonstrated better performance, achieving a Spearman correlation coefficient greater than 0.5 in 5 out of 11 systems, while ProteinMPNN is 0, validating its effectiveness in protein complex modeling. We evaluated the relationship between model-predicted scores and binding free energy for 1,877 single-point mutations and 675 multi-point mutations across 11 proteins (Fig. 5e). The overall Spearman correlation coefficient was 0.46.

Our pretrained model does not rely on ESM2 features, it focuses on understanding the backbone geometric structure and modeling amino acid types. These results suggest that the model effectively captures the relationships between sequence, structure, and function. This not only highlights its potential as a useful tool for protein engineering but also paves the way for further research and practical applications.

### Ablation Study of ProDualNet Enhancements

We trained and evaluated several ablation models to assess the efficacy of our proposed methods. First, we compared the performance of our pre-trained model, ProDualNet_single, on the protein sequence recovery. ProDualNet_single either outperforms or matches ProteinMPNN across three protein types (monomers, homologous oligomers, and heterooligomers) (Fig. 5f). This demonstrates the effectiveness of our model, which combines a heterogeneous graph network composed of both a message-passing neural network (MPNN)^19^ and Transformer layers^45^, enabling it to capture the relationship between protein sequences and structures with greater precision. We also tested the performance of ProteinMPNN after fine-tuning for dual-target protein design, further demonstrating the advantages of ProDualNet (Fig. 5g).

To validate the impact of our NoiseMix strategy, we designed an experiment to assess its influence on dual-target protein design task modeling without recycling. By mixing noise-augmented data with real dual-target data at varying ratios, we found that models trained with the augmented data (recovery = 0.581) significantly outperformed those trained solely on raw data (recovery = 0.546) or using a training method that does not sample by protein clusters (recovery = 0.532) (Fig. 5h). This result underscores the positive effect of noise augmentation on dual-target sequence design performance. In addition, we present the training and validation loss curves during the fine-tuning process, see supplementary materials. When the NoiseMix strategy is not applied, the other two fine-tuning methods lead to severe overfitting, which further highlights the effectiveness of the NoiseMix strategy (Fig. S1).

Next, we conducted a detailed evaluation of the model’s performance on the training dataset and test set (Fig. 5g). The experimental results showed that the model using the noise-augmentation strategy exhibited consistent performance between the training and testing datasets, and the recycle strategy is equally effective for the training dataset. Furthermore, ProDualNet still outperforms the ProteinMPNN_mean fine-tuned with NoiseMix in terms of sequence recovery, with a score of 0.581 compared to 0.534.

Furthermore, we validated the effectiveness of both the recycling strategy and ESM2 feature embedding (Fig. 5i). In our experiment, we varied the recycling parameter to assess its impact on model performance. Setting the recycling parameter to 1 yielded the best results in terms of both recovery and runtime efficiency. Specifically, this setting significantly outperformed models without recycling and those using only ESM2 pre-trained features. Moreover, the performance with a recycling parameter of 1 was comparable to that with a setting of 2, but with a significantly reduced computational time. These findings suggest that appropriately recycling intermediate results can effectively enhance the sequence design capability. In addition, when ProDualNet was trained without using ESM2 features, it achieved a sequence recovery of 0.54, which still outperformed ProteinMPNN_mean’s 0.52.

## Discussion

In this study, we present ProDualNet, a sequence design model for dual-target protein design tasks, aimed at more comprehensively modeling the intricate interactions between multi-target proteins. This model extends protein complex design from single-target protein (e.g., ProteinMPNN^19^, ProBID-Net^25^) to dual-target protein sequence design and addresses the scarcity of dual-target protein experimental data by employing a noise-augmentation approach (NoiseMix). This approach effectively mitigates the limitations of dual-target protein experimental structures, enhancing the model’s performance and stability in dual-target protein complex design tasks.

We evaluated the model in various scenarios, and ProDualNet demonstrated significant advantages in dual-target protein design tasks. Compared to the multi-state design method ProteinMPNN_mean^24^, ProDualNet showed significant improvements in total residue recovery, interface residue recovery, and perplexity. Additionally, by using structure prediction models like AlphaFold3^29^, ProDualNet outperformed the comparison methods in terms of PAE, PLDDT, and ipTM overall scores. Notably, when designing proteins that interact with different targets, ProDualNet better handles the conformational changes between different protein receptors and the variations in binding interfaces, improving design quality and success rate.

The study of amino acid site changes’ effects on protein function is crucial for understanding the relationship between protein sequence, structure, and function^27,36,46,47^. The ProDualNet model proposed in this study performs excellently in dual-target peptide drug screening predictions. Through evaluation on the dual-target dataset based on GCGR and GLP-1R, ProDualNet significantly outperforms traditional methods (such as AlphaFold3) in top-K% screening based on binding affinity, with a time advantage. Furthermore, ProDualNet demonstrates outstanding performance in mutation effect prediction. This advantage suggests that design models like ProDualNet are comparable to traditional large protein language models in terms of understanding sequences, structures, and functions.

Although this study is primarily based on in silico validation, we have proven the effectiveness and reliability of this method in dual-target protein design. Future research will further validate ProDualNet’s performance in practical tasks through wet lab experiments and extend it to design scenarios involving more complex proteins and other biomolecular interactions, such as RNA, DNA, metal ions, small molecules, etc^11,16^.

## Materials and Methods

### Pretraining Dataset

To construct the dual-target protein dataset, we first curated our dataset from PDBbank. Building on the methodology of ProteinMPNN^19^, we clustered proteins in the database into 25,361 clusters based on a 30% sequence identity cutoff using mmseqs2^48^. From these, we selected 766 protein clusters known to interact with multiple targets. The dataset was then split into 466 clusters for training and 300 clusters for testing (Fig. 1c).

For pretraining, we utilized 20,137 clusters from the database as the training set, with 1,662 clusters allocated for testing and 1,662 clusters for validation. It is important to note that the clusters in the multi-target test set are entirely derived from the pretraining validation and test datasets, ensuring that there is no data leakage.

### Fine-Tuning Dataset and Test Dataset

Following pretraining, the fine-tuning stage involved extracting 300 dual-target protein clusters from the testing and validation sets. These were used for the dual-target design task. The remaining 466 multi-target protein clusters from the training set were utilized for fine-tuning, focusing on dual-target protein interactions. To address the scarcity of training data, we employed a NoiseMix strategy. In each fine-tuning epoch, the 466 multi-target protein clusters from the training set were combined with random protein samples from the training dataset. These samples were augmented with Gaussian noise before being fed into the model for training. The detailed training procedure is provided in the supplementary materials (Table S2).

To evaluate the model’s robustness under various conditions, we established three test sets based on different criteria, such as sequence similarity, conformational changes, and binding interfaces. These test sets were derived from the same 300 protein clusters through stratified sampling, and some overlap may occur between them:

- **Test Set 1:** This set consists of 159 dual-target protein complex pairs, with homologous similarity between the two target proteins below 30%.
- **Test Set 2:** This set contains 52 protein complex pairs, where the RMSD of conformational changes between protein-protein interactions exceeds 2 Å, and the similarity between the two receptors is below 50%.
- **Test Set 3:** This set includes 38 protein complex pairs, where the binding interfaces of the designed protein do not overlap with either of the two target proteins and receptor similarity is below 30%.

These three test sets were designed to evaluate the model’s performance under varying conditions of target protein similarity, conformational change, and binding specificity.

### GCGR-GLP1R Dual-target Dataset

To assess the performance of ProDualNet in dual-target peptide drug design, we conducted a benchmark evaluation using a dual-target agonist dataset based on GCGR and GLP-1R. The dataset comprises 46 protein sequences, each 29 amino acids in length, with experimentally determined EC50 values for both GCGR and GLP-1R receptors^37^. Importantly, each sequence differs by at least one amino acid, with an average variation of 14 amino acids, rendering this task more complex than single mutation predictions. Notably, all test set sequences were excluded from the model’s training data, ensuring a robust and unbiased evaluation.

### Fitness Dataset for Mutation Prediction

For the assessment of thermostability mutations, we used two databases, ProteinGym DMS^27^ and DMS_stability^39,40^. From ProteinGym DMS, we selected 65 proteins with mutations related to thermostability, consisting of 67,636 single-point mutations and 65,579 higher-order mutations. For the DMS_stability dataset, we used their processed data, which includes 298 proteins and a total of 271,231 single-point mutations.

Additionally, to evaluate the performance in predicting the fitness of complex mutations, we selected mutation data from the SKEMPI V2 database^44^. This subset includes antibody mutations with more than 100 entries, comprising 11 proteins, 1,877 single-point mutations, and 675 higher-order mutations.

These diverse mutation datasets provide a robust benchmark for evaluating the ability to predict the stability of mutations within both individual proteins and protein complexes.

### Protein Sequence Design Overview

Unlike many existing structure-based sequence design models, which predict amino acid identities as conditional probabilities based on the protein backbone structure^15,16^, the protein complex design model predicts sequences by incorporating the relationship between the protein complex’s backbone structure and the target receptor sequence information^19,25^. For the dual-target protein design task, the model needs to consider the combination of two different targets information at the same time.

Protein sequence design tasks can be categorized based on the extent of structural information used. In the protein inverse folding problem, the model is tasked with predicting a protein sequence 𝑆_pre_ that corresponds to a given protein backbone structure. The prediction is formulated as^19,20^:

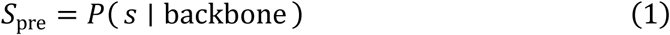

where 𝑠 represents the protein sequence, and the backbone refers to the structural scaffold of the protein.

When the protein sequence design task is extended to protein complex design, the challenge shifts from designing a single protein sequence to considering the interactions between proteins. The designed sequence must not only match the backbone structure of the protein complex but also interact with a specific receptor sequence.

The formulation for this case is:

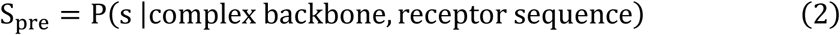

where the complex backbone refers to the structural arrangement of the entire protein complex, and the receptor sequence represents the sequence of the target protein or binding site with which the designed sequence must interact.

For dual-target protein design, the complexity increases further, as the designed protein interacts with two target proteins, forming two distinct hetero-oligomeric structures. Therefore, the model must take into account the interactions with both targets simultaneously to effectively sample the design space. The predicted sequence 𝑆_pre_ is now formulated as:

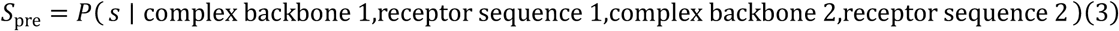

where the model processes the structural and sequence information from two different protein complex backbones and their respective receptor sequences to design a sequence that can simultaneously bind to both targets.

The existing methods simplify the joint distribution above into a probability averaging approach, as shown in the following formula^19,20^:

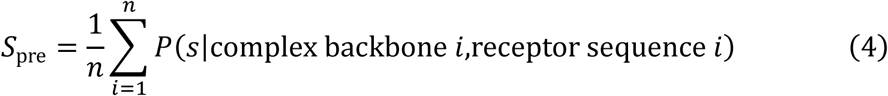

### ProDualNet Architecture

The architecture for the dual-target protein design task involves a shared encoder-decoder framework, where the encoder processes the structural and sequence information of both targets. This encoder, composed of four layers of heterogeneous graph networks, captures the relationships between the two target structures through a unified embedding space. Each layer of the heterogeneous graph network consists of an MPNN (Message Passing Neural Network) and a Transformer network. The encoder learns to capture the interdependencies between the two multi-target hetero-oligomeric structures through pretraining followed by fine-tuning.

To capture these interdependencies, for each layer of the heterogeneous graph network, the feature vectors 𝑧_i,1_ and 𝑧_i,2_ corresponding to the two different conformations of the designed protein complex are combined via a weighted sum:

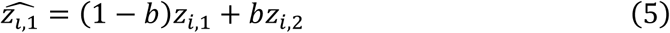

where 𝑏 = 0.2 is a hyperparameter that controls the contribution of each target structure to the final representation. This weighted combination allows the model to learn shared features that are relevant to both targets.

To further model the joint sequence distribution for both targets, the softmax function is applied to the combined decoder feature representation, enabling efficient model optimization:

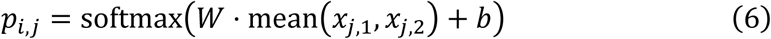

Where 𝑊 is the weight matrix, and 𝑏 is the bias term. The vectors 𝑥_j,1_and 𝑥_j,2_ represent the feature vectors of the two distinct target conformations. The value 𝑝_i,j_represents the predicted amino acid type at position 𝑖, based on the feature vector 𝑥_j_ at position 𝑗 of the protein. In this case, 𝑖 = 𝑗.

The mean operation ensures that the features from both targets are effectively fused before prediction. This approach allows the model to account for the structural and sequence dependencies between the two targets, facilitating the design of sequences that can interact with both targets simultaneously.

### NoiseMix Strategy

Due to the limited availability of high-quality dual-target protein data, directly pretraining and fine-tuning on this data may lead to overfitting. To address this, we adopt a two-phase training strategy consisting of pretraining followed by fine-tuning. In the pretraining phase, the model is trained on a larger, more general dataset that includes single-target protein sequences or protein complexes with less specific receptor information, similar to the strategy used in ProteinMPNN. This phase aims to teach the model the fundamental mappings between protein structure and sequence, and to understand potential functional relationships. We term this model ProDualNet_single. Specific details of the dataset and procedure are shown in Figure 1(c), where we avoid any possible data leakage through protein family clustering.

Following pretraining, the model is fine-tuned on a smaller, domain-specific dual-target dataset that incorporates a noise augmentation strategy, which we refer to as NoiseMix. This strategy involves blending noise-augmented data, generated by modifying the structures of non-dual-target proteins, with real dual-target experimental data during the fine-tuning process. The amount of noise-augmented data used is treated as a hyperparameter, allowing it to be adjusted for optimal training performance. This approach enhances the model to generalize and reduces the risk of overfitting. The effectiveness of this noise-enhanced fine-tuning strategy in improving ProDualNet’s performance on complex dual-target design tasks is demonstrated in the ablation study.

### Noise Augmentation

During the fine-tuning phase, to leverage more available training data and further improve generalization, we introduce a noise augmentation strategy. In this approach, Gaussian noise is added to the coordinates of non-dual-target protein complexes 𝑟_i_ to generate synthetic structure pairs (𝑟_i,1_, 𝑟_i,2_), where 𝑟_i,1_ and 𝑟_i,2_ are noisy versions of the same complex.

Mathematically, this process is represented as:

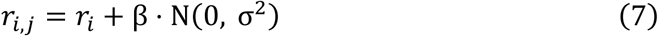

where N(0, σ^2^) denotes Gaussian noise with mean 0 and variance σ^2^, and β is a scaling factor that controls the magnitude of the noise (e.g., β=0.2, σ^2^=1). These noisy structure pairs are incorporated into the training process, augmenting the available dual-target data and enhancing the model to generalize across a broader variety of protein conformations.

By training on both real and synthetic data, the model is exposed to a richer set of protein structures, which significantly improves its ability to handle the complexity of dual-target interactions. This combined training approach increases the robustness of the model, enabling it to design sequences that can interact with both targets in a variety of configurations.

### Loss Function

The loss function used to train the model measures the discrepancy between the predicted and true amino acid types at each position in the protein sequence. For a sequence of length 𝑙, where each position 𝑖 is represented by a feature vector 𝑥_i_ the model predicts the probability distribution over the 20 amino acid types using the softmax function:

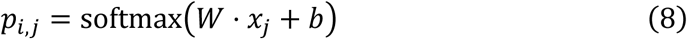

where 𝑊 and 𝑏 are the weight matrix and bias term, respectively. 𝑝_i,j_ represents the prediction of the amino acid type at position 𝑖 based on the feature 𝑥_j_ at position 𝑗 of the protein.

The total loss is the weighted sum of cross-entropy losses for each position 𝑖, as well as for its neighboring positions 𝑖 − 1 and 𝑖 + 1. The individual cross-entropy loss at position 𝑖 is defined as:

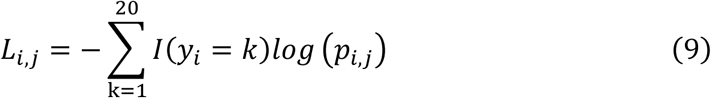

Where 𝐼(𝑦_i_ = 𝑘) is the indicator function that equals 1 if the true amino acid at position 𝑖 is 𝑘, and 0 otherwise. The overall loss is computed as:

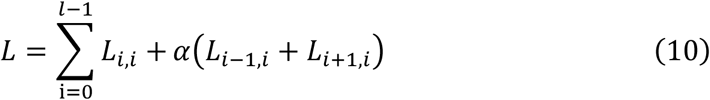

where 𝛼 is a hyperparameter controlling the weight of neighboring positions in the loss calculation.

### Training Details

**Pretraining:** For training, we used a single NVIDIA A100-SXM4-40GB GPU. The model is pretrained using the Adam optimizer with parameters β_1_ = 0.9, β_2_ = 0.98 and ɛ = 10^-9^, following the learning rate schedule of ProteinMPNN. The pretraining phase lasts for 5,000K steps, with α set to 0.25. The training dataset consists of 20,137 protein clusters.

**Fine-tuning:** Following pretraining, the model undergoes fine-tuning using the Adam optimizer with a learning rate of 0.00001 for an additional 20K steps. This phase focuses on optimizing the model for dual-target protein interactions, with α=0. For models utilizing ESM2 protein language model features, we fill the target sequence features with a zero tensor and apply a recycled training approach. Specific training details are provided in the supplementary materials. During fine-tuning, each epoch includes a mixture of dual-target data from 466 protein clusters with multi-target interactions, as well as 4,000 protein clusters randomly sampled from the pretraining dataset, augmented with noise, the detailed training algorithms are provided in Algorithm 1 and Algorithm 2 in the supplementary materials.

## Data and Code availability

Data and code are available at https://github.com/chengliu97/ProDualNet.

## Acknowledgements

This study was supported by National Natural Science Foundation of China (ID: 12171318), Shanghai Science and Technology Commission (ID: 24JS2810200,21ZR1436300,23XD1401900,23DZ2290600), Medical Engineering Cross Fund of Shanghai Jiao Tong University (ID: YG2023ZD21). The National Key Research and Development Program of China (2020YFA0907700 and 2023YFF1205102), the National Natural Science Foundation of China (21977068 and 32171242), and the Fundamental Research Funds for the Central Universities (YG2023LC03). The computations in this paper were run on the Siyuan-1 cluster supported by the Center for High Performance Computing at Shanghai Jiao Tong University.

